# Biochemical characterization of Pectin Methylesterase Inhibitor 3 from *Arabidopsis thaliana*

**DOI:** 10.1101/2022.04.14.488319

**Authors:** Fan Xu, Martine Gonneau, Elvina Faucher, Olivier Habrylo, Valérie Lefebvre, Jean-Marc Domon, Marjolaine Martin, Fabien Sénéchal, Alexis Peaucelle, Jérôme Pelloux, Herman Höfte

**Affiliations:** Université Paris-Saclay, INRAE, AgroParisTech, Institut Jean-Pierre Bourgin (IJPB), 78000, Versailles, France; UMRT INRAE 1158 BioEcoAgro – BIOPI Biologie des Plantes et Innovation, SFR Condorcet FR CNRS 3417, Université de Picardie, 33 Rue St Leu, 80039 Amiens, France

## Abstract

The *Arabidopsis thaliana PECTIN METHYLESTERASE INHIBITOR 3 (PMEI3)* gene is frequently used as a tool to manipulate PME activity *in vivo*, in studies assessing the role of pectin de-methylesterification in the control of cell expansion. One limitation of these studies is that the exact biochemical activity of this protein has not yet been determined. In this manuscript we produced the protein in *Pichia pastoris* and characterized its activity *in vitro*. Like other PMEIs, PMEI3 inhibits PME activity in acidic pH conditions for a variety of cell wall extracts and for purified PME preparations, but doesn’t affect PME activity at neutral pH. This suggests that the previously observed *in vivo* effects reflect the inhibition of PME activity at low pH. The protein is remarkable heat stable and shows higher activity against PME3 than against PME2, illustrating how different members of the large PMEI family can differ in their specificities towards PME targets. Finally, application of purified PMEI3 on *Arabidopsis thaliana* seedlings showed a dose-dependent inhibition of homogalacturonan de-methylesterification and root growth. Purified recombinant PMEI3 is therefore a powerful tool to study the connection between pectin methylesterification and cell expansion.

## Introduction

A central question in plant biology is how growth can occur despite the presence of strong cell walls (Cosgrove, 2005). Cell walls of growing cells (i.e. primary cell walls) are highly dynamic polymer assemblies primarily consisting of cellulose and the matrix polysaccharides hemicelluloses and pectins. Pectins are galacturonic acid (GalA) - containing polysaccharides among which the linear polymer homogalacturonan (HG) is the most abundant matrix polysaccharide, for instance representing 20% of the total cell wall polysaccharides, in *Arabidopsis thaliana* leaves (Zablackis et al., 1995) or even 50% in onion epidermis cells (Wilson et al., 2021). *In vivo* studies show that the metabolism of HG plays a critical role in the control of cell expansion and plant morphogenesis (Andres-Robin et al., 2018; K. T. Haas et al., 2020; Jobert et al., 2021; Peaucelle et al., 2015, 2008; Phyo et al., 2017; Qi et al., 2017; Stefanowicz et al., 2021; Wachsman et al., 2020) but the underlying mechanisms are still poorly understood.

HG is synthesized in a highly methylesterified form in Golgi bodies and delivered to the surface via secretory vesicles. In the cell wall, de-methylesterification by pectin methylesterases (PMEs, EC 3.1.1.11) uncovers the negatively charged carboxylic acid groups, which dramatically changes the physical properties of the individual polymers and polymer assemblies (Atmodjo et al., 2013; Haas et al., 2021). In addition, depending on the methylesterification pattern, HG is more or less sensitive to enzymatic degradation by polygalacturonases and pectate lyases (Hocq et al., 2017). PMEs are part of large gene families (66 members in *A. thaliana* (Wolf et al., 2009)), with different expression patterns, but also different biochemical characteristics. For instance, most plant PMEs have processive activity, generating blockwise de-methylesterified HG, in contrast to characterized fungal PMEs, which are non-processive, generating more randomly de-methylesterified HG (Wolf et al., 2009). PME activity is also strongly pH dependent, with most PMEs showing neutral or alkaline pH optima. Interestingly, the pH also influences the mode of action of the enzymes. For instance, citrus PME and *Arabidopsis thaliana* PME2 are highly processive at pH 8 but not at pH 5 (Hocq et al., 2021). The biological significance of this behavior, in particular in relation to growth control remains to be determined. PME activity is also regulated by PME inhibitors, which are also encoded by large gene families in vascular plants (76 in *A. thaliana* (Wolf et al., 2009)). PME activity was shown to have a critical role in the control of cell expansion and morphogenesis. The inducible overexpression of PME5 and PMEI3 promoted and inhibited cell wall expansion respectively, and both treatments caused the loss of growth anisotropy (Peaucelle et al., 2015). PMEI3 overexpression also caused the inhibition of organ formation in the inflorescence meristem (Peaucelle et al., 2011). Similar observations were made in which PME5 and PMEI3 overexpression respectively caused an increased and reduced gynoecium length (Andres-Robin et al., 2018), or reduced or increased cell expansion and loss of organ asymmetry in *A. thaliana* leaves (Qi et al., 2017). Finally, PMEI3 overexpression also led to the Inhibition of lateral root formation (Wachsman et al., 2020).

Surprisingly, some of these studies revealed opposing effects of changing the degree of HG methylesterification on cell wall mechanics and expansion rate. Indeed, AFM studies showed that, depending on the context, demethylesterification caused an increase or a decrease in wall elasticity and growth rate (Peaucelle et al., 2015; Phyo et al., 2017). It has been proposed that these opposite effects may be related to the pH-dependent, processive or non-processive nature of certain PMEs (Hocq et al., 2017). In this view, processive activity generates stretches of acidic GalA, which can form cooperative Ca^2+^ crosslinks (the so-called egg boxes) and which would cause cell wall stiffening and growth inhibition. In contrast, randomly demethylated HG would form a more elastic Ca^2+^ cross-linked network (Vincent et al., 2013). An alternative view explaining PME-dependent growth was recently proposed based on the observation, using super-resolution optical microscopy, of oriented HG filaments in anticlinal cell walls of leaf pavement cells (Haas et al., 2020). In this view the de-methylesterification of HG leads to a volume increase of HG filaments, which would be a driver for cell expansion.

The above-mentioned studies used the inducible expression of the putative PMEI inhibitor PMEI3 (At5G20740) to manipulate pectin methylesterification. Despite its frequent use *in vivo*, the biochemical activity of PMEI3 has not been characterized so far. This knowledge is essential however, for the correct interpretation of the induced mechanical, cellular and macroscopic phenotypes. A few other PMEIs (PMEI4, 7 and 9) have been heterologously expressed and biochemically characterized (Hocq et al., 2016; Senechal et al., 2015). This showed, for different PMEIs, pH-dependent (PMEI4, 7) or pH-independent (PMEI9) ineractions with target PMEs. In this report, we produced the PMEI3 protein in *Pichia pastoris* and characterized its activity and specificity. We show a pH-dependent inhibition of PME activity. Interestingly the inhibitory activity and the pH dependence varied against different PME enzymes, showing the potential biochemical specialization of PME-PMEI pairs. We also show that purified PMEI3 is a potent tool for the *in vivo* manipulation of PME activity to study its role in cell expansion.

## Results

### Expression in Pichia pastoris and purification of PMEI3

A codon-usage-optimized sequence of *Arabidopsis thaliana* PMEI3 (At5G20740), fused at its N-terminus to a secretion signal peptide and at its C-terminus to 6XHIS and cMYC tags, was expressed in *Pichia pastoris* X33 (Figure 1A). The mature protein could be detected in the culture supernatant using immunoblotting with anti-HIS antibodies (Figure 1B). Upon His-tag affinity purification (Figure S2A), a band of the expected molecular weight (∼25kDa) was observed on Coomassie-stained SDS PAGE (Figure 1C). The identification of the PMEI3 protein was verified using immunoblotting with anti-HIS antibodies (Figure 1C) and LC-MS/MS on the band extracted from the gel, which detected in the tryptic digest 4 peptides matching PMEI3 (Figure 1D).

**Figure 1:**
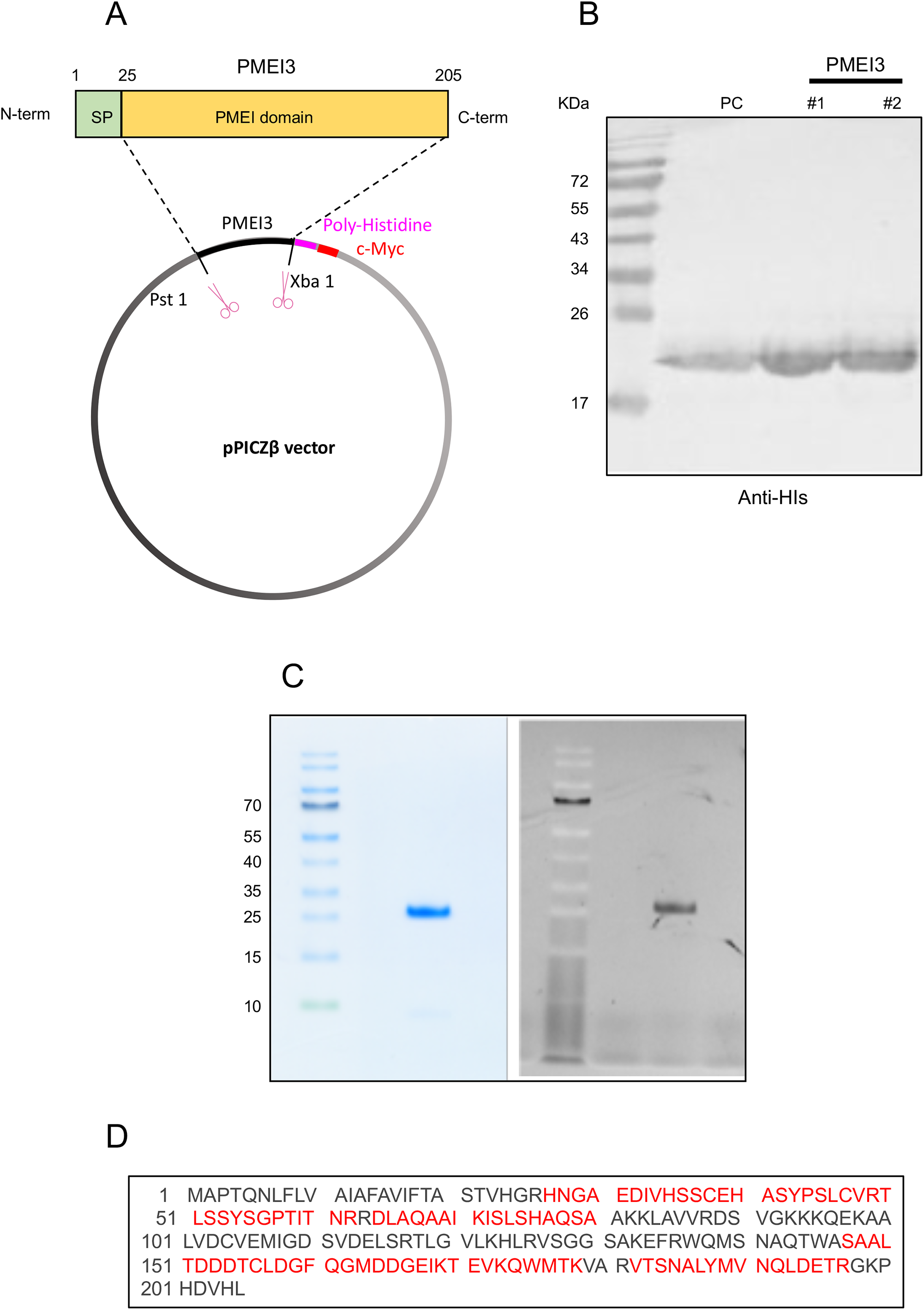
Expression in Pichia pastoris and purification of PMEI3. A. Scheme of the expression construct, B. Immunoblot with anti-HIS antibodies of 2 culture supernatants (#1 and #2) and positive control (PC) of purified PMEI3. C. Coomassie stained gel (left) anti-HIS immunoblot (right) of purified protein. D. PMEI3-matching tryptic fragments (in red) from the purified protein identified by MS-MS.

### pH-dependent inhibition of PME activities

We next characterized the inhibitory activity of PMEI3 on a number of PME preparations. The PME activity was first measured on commercial citrus pectin (Degree of Methylesterification (DM) 85%) at pH 7.5 using a colorimetric assay for methanol production on dilutions of the enzyme preparation (Figure S1). Non-saturating dilutions were chosen for the PMEI assays (1/100 dilution for Orange PME and 1/10 dilution for flower and root PME preparations). The activity of purified PMEI3 was assayed using a gel diffusion assay at pH 5, 6.3 and 7.5 respectively. The activity was quantified by measuring the radius of the red halo (Figure S2). We first tested the inhibiting activity of PMEI3 against a commercial PME preparation from orange peels. The results showed strong inhibition at pH 5 and decreasing inhibition from pH 6.3 to 7.5 respectively (Figure 2A-C). Surprisingly, boiling the sample did not (at pH 5) or only slightly (at pH 6.3) reduce the PMEI3 activity. This indicates that this inhibitor is able to correctly refold upon heat denaturation. A similar pH-dependent inhibitory pattern was observed against PME preparations from roots (Figure 2D-F) and inflorescences (Figure 2G-I) respectively. Interestingly, the inhibitory activity was stronger on the extracts from roots than from flowers at both pH 5 and 6.3. In conclusion, PMEI3 showed a pH-dependent inhibitory activity against one or more PME isoforms that are expressed in roots and flowers. The lower activity on flower extracts suggests that different PME isoforms may vary in their sensitivity to PMEI3 inhibition.

**Figure 2:**
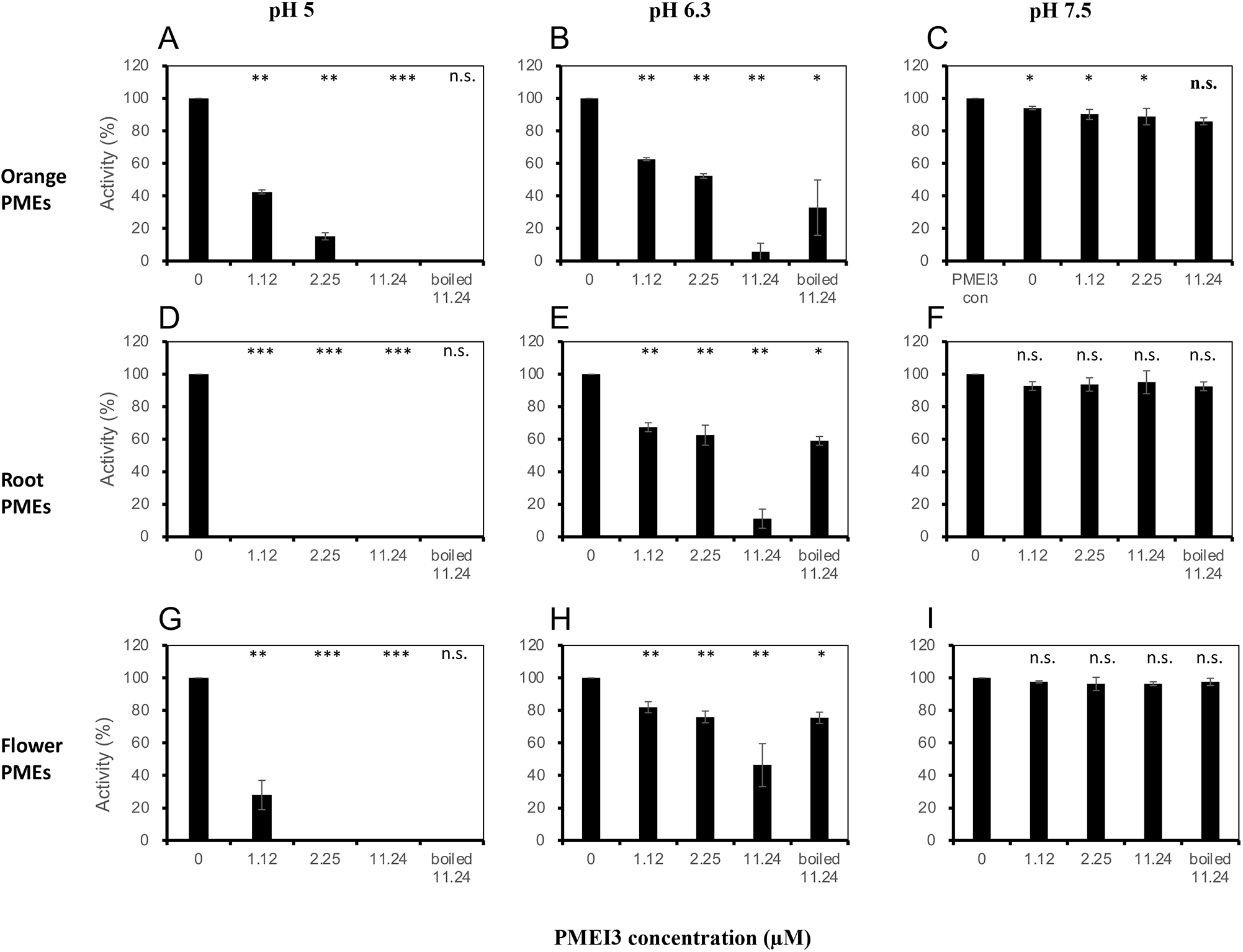
pH-sensitive inhibition of PME activity by PMEI3. Relative PME activity measured at pH 5 (A,D,G); 6.3 (B,E,H) and 7.5 (CF,I) in the presence of 0; 1.12 μM; 2.25 μM; 11.24 μM and boiled 11.24 μM PMEI3 in PME preparations from orange (A-C), *Arabidopsis thaliana* roots (D-F) or flowers (G-I). PME activity is expressed as % of the control without PMEI3. Error bars are SD (n=3) and stars refer to the P-value of the t-tests on the comparison with the 0 control or for the boiled samples for the comparison with non-boiled 11.24 μM samples. *:<0.05, **:<0.01, ***:<0.001, ns : not significant.

### Inhibitory activity and pH dependence of PMEI3 varies for different PME isoforms

To further investigate the specificity of PMEI3, we compared its activity against two heterologously expressed PME isoforms PME2 (At1g53830)(Hocq et al., 2021) and PME3 (At3g14310)(Guenin et al., 2011) (Figure 3). Both PME2 and PME3 are closely related isoforms with a broad expression pattern. PME3 has a broad expression pattern in roots and leaves (Figure S3). Based on mutant analysis, PME3 represented around 30% of the PME activity in root extracts (Guenin et al., 2011). PME2 instead, has a more restricted expression pattern in roots and flower petals (Figure S4). In addition, mutant analysis showed that the contribution of PME2 to the total PME activity was only marginal at least in the organ investigated (dark-grown hypocotyl) (Hocq et al., 2021). The latter may reflect the epidermis-specific expression pattern of this gene, suggested by the cell type-specific RNA expression data (Figure S5). PME2 was produced in *Pichia pastoris* and purified using ion exchange chromatography (Hocq et al., 2021). PME3 fused to a C-terminal 6xHIS-tag was transiently expressed in tobacco leaves and affinity purified (Senechal et al., 2015). Again, at pH 7.5, no PME inhibitory activity was observed against either one of the PMEs, with for the lower PMEI3 concentrations even a small but significant (t-test, P value = 0.008) PME3 promoting activity. Interestingly however, at pH 5, PMEI3 was at least 10-fold more active against PME3 than against PME2 (PME3 inhibition with 1.12 μM PMEI3 was comparable to PME2 inhibition with 11.2 μM PMEI3). In addition, at pH 6.3, PMEI3 was still active against PME3 but not against PME2. Together, these observations illustrate the preference of this PMEI isoform for PME3 relative to PME2.

**Figure 3:**
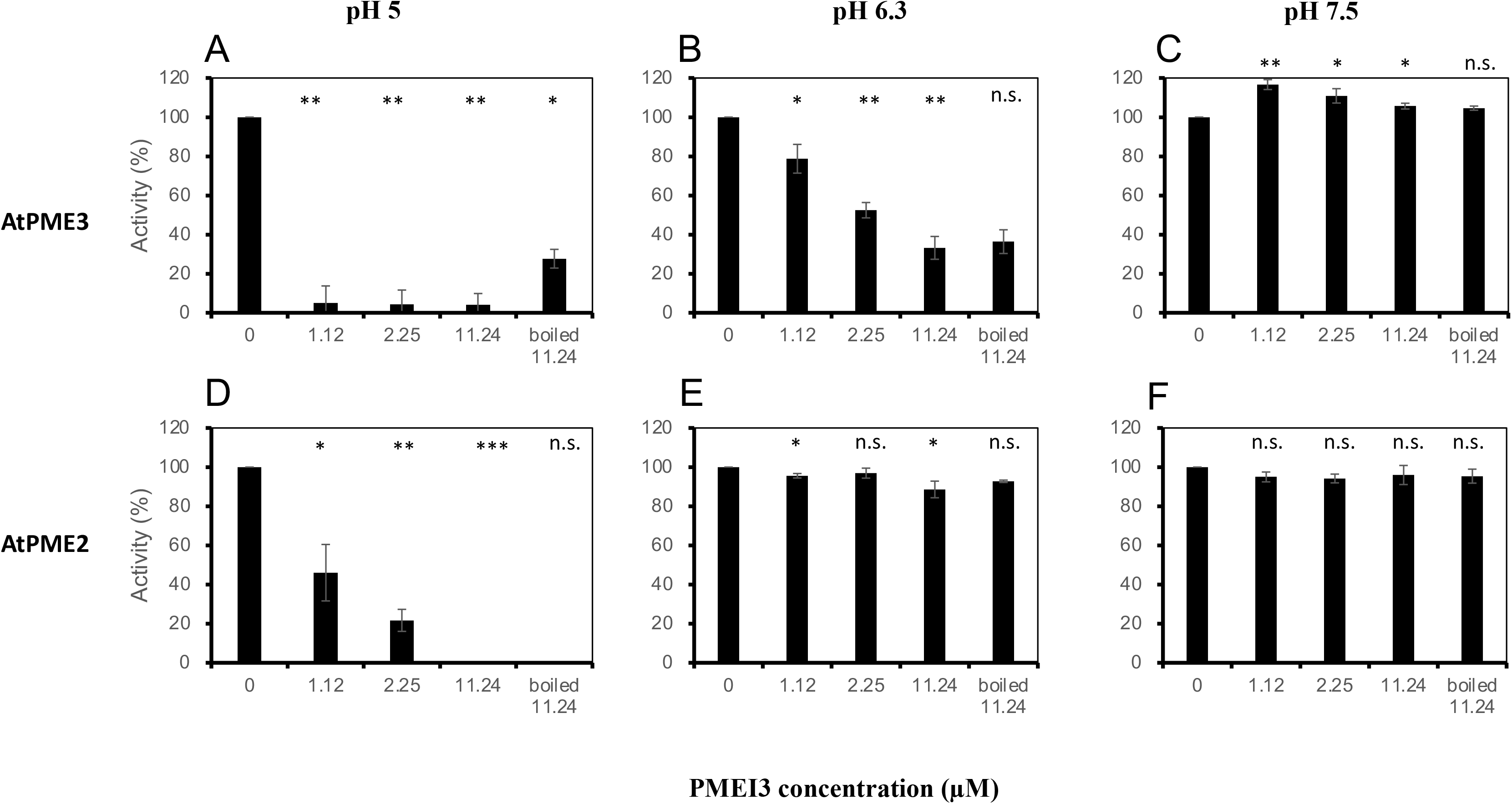
Specific PMEI3 activity against purified PMEs. Relative PME activity measured in purified PME3 (A-C) and PME2 (D-F) preparations. at pH 5 (A-D); 6.3 (B-E) and 7.5 (C-F) in the presence of 0; 1.12 μM; 2.25 μM; 11.24 μM and boiled 11.24 μM PMEI3. PME activity is expressed as % of the control without PMEI3. Error bars are standard deviations (n=3) and stars refer to the P-value of the t-tests on the comparison with the 0 control or for the boiled samples for the comparison with non-boiled 11.24 μM samples. *:<0.05, **:<0.01, ***:<0.001, ns : not significant.

### In vivo effect of PME inhibition

We next assessed the *in vivo* effect of the inhibition of PME activity by supplementing purified PMEI3 to the growth medium of *A. thaliana* seedlings. The small molecule PME inhibitor epigallocatechin-3-gallate (EGCG) was used as a positive control. EGCG caused a dose-dependent reduction in the root length of 4-day-old seedlings, relative to the mock-treated control, with an IC^50^ of around 55 μM (Figure S6). Similarly, the root length of 3-day-old seedlings grown in the presence of PMEI3 was strongly reduced in a dose-dependent fashion (Figure 4A,B) with an IC^50^ of 0.16 μM. The labeling of de-methylesterified HG with fluorescent COS^647^ (Mravec et al., 2014), showed a strongly reduced labeling in roots treated with 0.2 or 0.6μM PMEI3 relative to mock-treated controls (Figure 4C,D). Increasing the sensitivity of the detection also showed a dose responsiveness of the fluorescent signal (0.2 sat and 0.6 sat in Figure 4C,D). In summary, these results show that external supply of PMEI3, like the PME inhibitor EGCG, mimic the inhibitory effect on cell expansion of the inducible overexpression of the *PMEI3* gene (Peaucelle et al., 2015).

**Figure 4:**
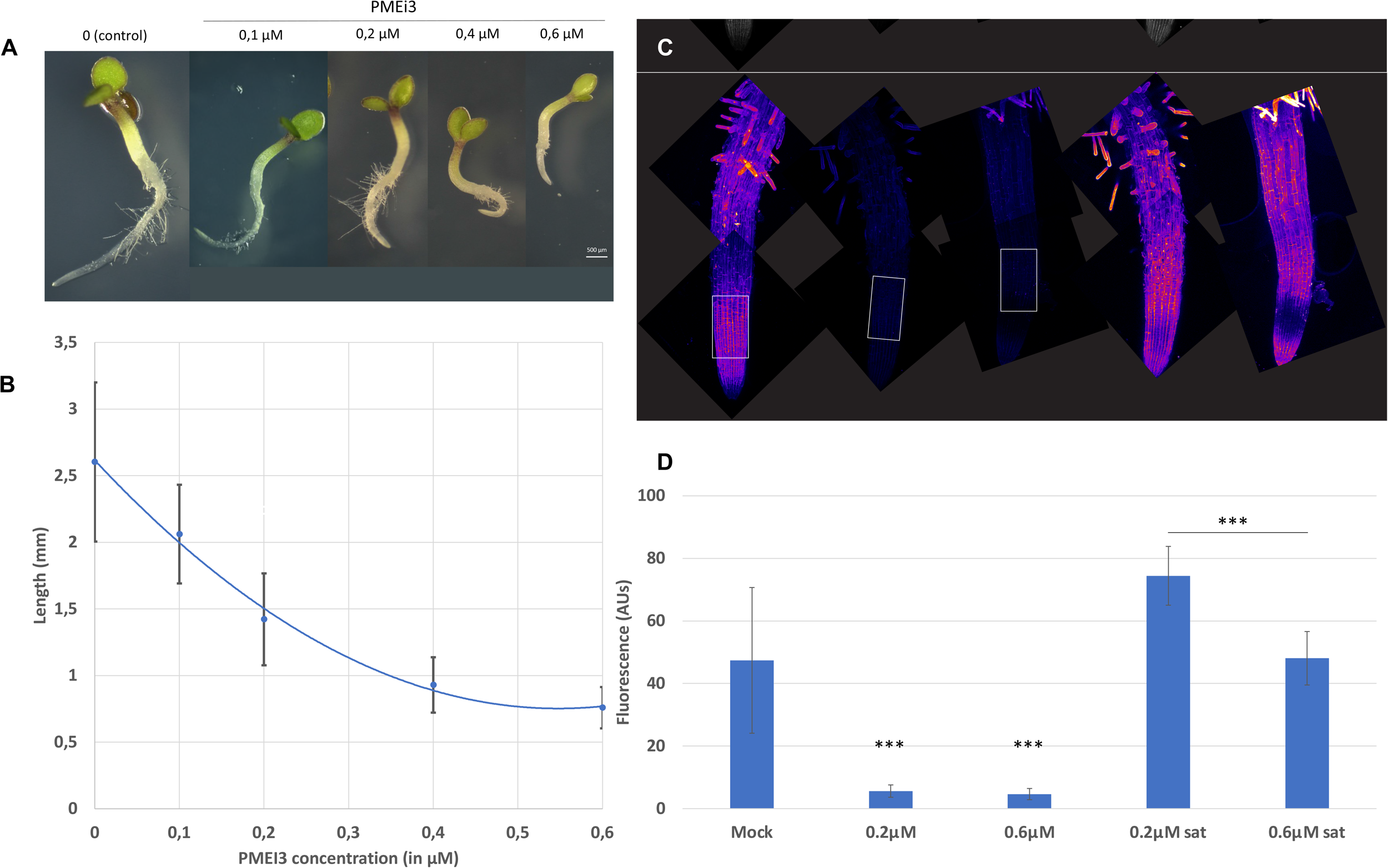
*In vivo* activity of purified PMEI3. A,B: Representative photographs (A) and primary root lengths (B) of 3-day-old *Arabidopsis thaliana seedlings*. Curve is a second degree polynomal fit (y = 3,3692x^2^ - 3,6899x + 1,007; R^2^ = 0,9947); IC50 = 0.16 μM. C,D: Representative photographs (C) and fluorescence quantification (D) of COS^647^-labeled roots. 0.2μM sat and 0.6μM sat refer to the same samples as for 0.2μM and 0.6μM but with an increased detection sensitivity. Two-day-old seedlings were transferred to 0; 0.2 μM or 0.63 μM PMEI3 and incubated for 24h before analysis. Error bars are SD (n > 30 for B and n=5 for D). *** t-test, P-value<0.001.

## Discussion

Heterologously-expressed PMEI3 inhibited the activity of a variety of PME preparations. Interestingly, PMEI3 showed quantitative differences in its inhibitory activity, with higher activity against root PMEs and PME3 relative to flower and orange PMEs and PME2. PME3 is the major PME activity in vegetative tissues including roots, whereas PME2 transcripts are more abundant in flower petals. This may explain the higher sensitivity of root PMEs relative to flower PMEs for PMEI3. The comparison of the closely related 3D structures of PME2 and PME3 may provide valuable information on the structural basis for these differences in specificity.

*In vitro* PMEI3 activity was strongly pH-dependent, with the highest inhibition at pH 5.5 and almost no or no inhibition at pH 7.5. Similar observations were made for PMEI4 and PMEI6 but not PMEI9, which also inhibited PME activity at pH 7.5 (Hocq et al., 2016). PMEI3, like PMEI4 and 6, is therefore expected not to totally prevent PME activity but only to sharpen its pH optimum *in vivo*. Interestingly, the pH was recently shown to also affect the de-methylesterification patterns at least for orange PME and *Arabidopsis thaliana* PME2 (Hocq et al., 2021). Indeed, whereas both enzymes were processive at pH 7.5, generating blockwise demethylated HG, their activity became non-processive at pH 5.5. This suggests that the strong effect of PMEI3 overexpression on growth and development (Andres-Robin et al., 2018; Peaucelle et al., 2015, 2011; Qi et al., 2017; Wachsman et al., 2020) might be related to a change in the HG methylation patterns, rather than the quantitative inhibition of PME activity. This is surprising given the strong overall reduction of COS^647^ staining of PMEI3-treated seedling roots, given the fact that COS^647^ selectively binds blockwise de-methylesterified HG (Figure 4C,D). To explain this, it is possible that the activity in the context of an intact cell wall does not reflect the *in vitro* specificity on isolated wall extracts or purified PMEs. In this context it is interesting to note that the specificity of PME activity was shown to change when the enzyme was incorporated in pectin or pectin/cellulose gels (Bonnin et al., 2019; Vincent et al., 2013). Alternatively, non-processive PME activity at low pH might be an essential step to prime subsequent processive activity. It will be interesting to see what the effect is on *in vivo* pectin methylesterification and plant development of PMEI9 overexpression, which is expected to suppress PME activity irrespective of the pH, and to what extent HG populations with different methylation patterns can have distinct *in vivo* functions. A striking example of a HG methylesterification pattern with a specific *in vivo* function was recently reported for the external cell wall of seed coat epidermis cells (Francoz et al., 2019). Indeed, a specific pattern, recognized by antibody LM20, and requiring the presence of PMEI6, appeared to recruit a peroxidase (PRX36) to a specific location, thus generating a brittle zone in the external cell wall facilitating cell wall rupture and mucilage release at this location during seed hydration.

Finally, we observed that the external supply of PMEI3 to the growth medium inhibited the growth of seedlings in a concentration dependent way paralleling the PME inhibitory activity, corroborating the observations that PMEI3 overexpression inhibits cell wall expansion (Andres-Robin et al., 2018; Peaucelle et al., 2015, 2011; Qi et al., 2017; Wachsman et al., 2020). The purified protein will be a valuable tool to study the *in vivo* dynamics of pectin metabolism and the developmental consequences of the manipulation of pectin methylesterification.

In part based on the findings in this study, it will be interesting to further explore the biochemical characteristics of other members of the large PME and PMEI families in plants.

## Materials and Methods

### PMEI3 heterologous expression and purification

pPICZα B is a 3.6 kb vectors used to express and secrete recombinant proteins in *Pichia pastoris*. Recombinant proteins are expressed as fusions to an N-terminal peptide encoding the *Saccharomyces cerevisiae* α-factor secretion signal. The vector allows high-level, methanol-inducible expression of the gene of interest in *Pichia*, and can be used in *Pichia* strain X-33. A plant codon-optimized PMEI3 sequence (see supplementary data) was inserted into plasmid pPICZαB and transformed into *E. coli* strain TOP10. Transformants was selected on Low Salt LB plates containing 50 μg/mL Zeocin. Eight transformants for each were analyzed by restriction mapping or sequencing to confirm in-frame fusion of PMEI3 with the α-factor secretion signal and the C-terminal tag. The appropriate transformants were cultivated to collect more plasmid. Plasmids were purified using a DNA purification kit (NucleoBond PC 100/BAC kit, obtained from Machery-Nagel). The recombinant plasmids were purified and then linearized using the Pme1 restriction enzyme. Recombinant plasmids (at a concentration > 500 ng/μL) were transformed into *Pichia pastoris* strain X33. Transformed *Pichia* was plated onto YPDS medium containing 50 μg/mL of Zeocin.

Transformants were selected and then cultivated in BMGY medium. After 12-h incubation at 37°C with 250 rpm shaking, the OD^600^ of the culture was measured. The recombinant *Pichia* was transferred into BMMY medium in a bigger volume (Table 7.2) for a large-scale expression. The OD^600^ were measured every 24 h to investigate the *Pichia* growth rate. In the meantime, 100% methanol was added to the medium to keep its final concentration at 0.5% (v/v). To make sure the recombinant proteins expressed correctly, the concentrated supernatant of the culture was tested using SDS-PAGE and/or Immunoblotting (details see below).

After 4-d expression, *Pichia* cells were removed by centrifugation at 5000 rpm for 5 min. To the protein-containing supernatant was added PBS buffer 10X (Eurobio, GAUPBS00-01), 400 mM NaCl, 5 mM imidazole and 500 μL cOmplete Hig-tag purification Resin (obtained from Roche, ref 05893682001). The mixture was gently shaken at 4°C for 4 h in dark. The supernatant was removed by centrifugation at 8000 rpm for 5 min at 4°C. The resin was washed by 5 ml of wash buffer 1 (PBS 1X with 400 mM NaCl and 5 mM imidazole), 5 ml of wash buffer 2 (PBS 1X with 400 mM NaCl and 30 mM imidazole) and eluted by elution buffer (PBS 1X with 400 mM NaCl and 400 mM imidazole) for 3 times (800 μL for each). The 3 elutes were collected and centrifuged at 8000 rpm for 30 min by using 3-kDa filter (Amicon Ultra-2 mL, UFC200324). The buffer change was performed using PD SpinTrap G-25 column. Final protein concentration was measured with a Bradford assay. The purified PMEI3 was frozen in liquid nitrogen and stored at -80°C until use.

### Cell lysis and immunoblotting

*Pichia* cells were pelleted, rapidly frozen by liquid nitrogen and then stored at -80°C before use. Cells were resuspended in phosphate buffer pH 7.0 (2 mL for 1 mL pellet) containing 300 μl of protease inhibitor 1x (Constant systems Ltd. United Kingdom).

Proteins were resolved using a 15% acrylamide/bisacrylamide SDS-PAGE (Table 7.3), with a running buffer composed of 25 mM Tris-base, 192 mM glycine, and 0.1% SDS at pH 8.7 (for Figure 1B) or NuPage Bis/Tris 4-12 % acrylamide using MOPS running buffer (Figure 1C). 13 μL of protein was added to 15 μL Laemmli buffer 2X and 2 μL of 1 M imidazole. The protein mix was heated at 95°C for 20 min. 15 μL of protein mix was loaded into a well of a SDS-PAGE gel. After electrophoresis at 100 mA for 100 min, the gel was stained with Pageblue protein staining solution (Thermo Scientific, catalog no. 24620) and destained with distilled water for 6 h. After migration of proteins, the gel SDS-PAGE was immerged in cathode buffer (25 mM of Tris pH 9.4, 40mM of Glycine, 10% of ethanol) for 15 min. A piece of Hybond-P PVDF membrane (GE Healthcare, catalog no. RPN303F) was placed in 100% ethanol for 15 seconds and rinsed by water for 2 min. Then the membrane was incubated in anode buffer II (25 mM of Tris pH 10.4, 10% of ethanol) for 5 min. The membrane, gel, and 7 same-sized Whatmann papers (previously immerged in anode I, anode II and cathode buffers) were placed. The transfer “sandwich” was exposed to 25 V and 1 A for 30 mins by a Trans-Blot TURBO transfer system (Bio-Rad, catalog no. 170-4155). The membrane was incubated in 4.0% dry milk in PBS buffer, 0,1% Tween20 for 30 min and then incubated for 1 h at room temperature under shaking (50 rpm) in the same buffer containing 1/5000 of primary antibody raised against poly-His tag directly coupled with peroxidase (Sigma, catalog no. A7058). After washing 3x by PBS 1x, 10 mins for each, 500 μL of DAB peroxidase substrate was added (Thermo Scientific, catalog no. 34002). Imaging was performed with a LAS-4000 according to the user manual. The concentration of proteins was determined according to Bradford using a BSA standard (1976)

### Cell wall-enriched protein extraction

Fifty mg of frozen material of 10-day-old roots and 4-week-old flowers were ground with liquid nitrogen. 1 ml buffer (containing 50 mM Na^2^HPO^4^, 20 mM citric acid, 1 M NaCl, 0.01% Tween 20 at pH 7.0) was added. The solution was shaken at 250 rpm for 1 h. The extracts were clarified by centrifugation at 10000 rpm for 30 min at 4°C. The supernatant was filtered using an Amicon ultracentrifuge filter 0.5 ml/10000 MWCO (Millipore, catalog no. UFC5010BK) in order to remove salts. Protein concentration was determined by the Bradford method (details see below).

### Methanol colorimetric assay

Activity of AtPME2, AtPME3, citrus PME, and PMEs from flowers and roots were determined using a colorimetric microassay adapted from Klavons and Bennett (1986). The reaction solution contained 5 μL of purified protein, 85% methylesterified (DM85) Citrus pectin (Sigma P9561) at 1mg/ml, 0.008 U of *Pichia pastoris* alcohol oxidase (Sigma A2404) and 50 mM sodium phosphate buffer (pH = 7.5) to a final volume of 100 μl. AtPME2, AtPME3, and citrus PME were prepared at 3 concentrations (1, 1/10 and 1/100 of purified proteins) and PMEs from flowers and roots at 2 concentrations (1 and 1/10). The mixture was incubated at 28°C for 30 min. A first OD reading 420 nm was done on a BioTek PowerWaveXS2 spectrometer. 100 μL of developing solution (2 mM ammonium acetate, 0.02 M pentane-2,4-dione, 0.05 M glacial acetic acid) was added and the solution was incubated at 68°C for 15 min after which a second reading was done. A standard dilution series from 0 to 10 nmol of methanol was included in each batch. Results were expressed as nmol MeOH/min^-1^.μL^-1^ of protein using the methanol standard curve.

### Gel diffusion assay

DM85 Citrus pectin (Sigma P9561) was dissolved in 0.1 M citric acid and 0.2 M Na^2^HPO4 buffer at pH 5.0, 6.3 and 7.5 (Hocq et al. 2017; Sénéchal et al. 2015 and 2017; Downie et al. 1998). One percent (w/v) agarose was added to the pectin buffer and heated until the agarose had dissolved. The solution was cooled to 60°C and transferred to a Petri dish (50 ml per dish) and allowed to solidify. Wells were punched in the gel with a 2-mm-diameter plastic sucker and the gel plug removed by aspiration. The gel was nicked and the orientation of the nick marked on the outside of the Petri dish with a marker. Ten μL of enzyme mixture was pipetted into each well. Once loaded, the lids were placed on the dishes, the wells were incubated at 37°C for 16 h. After incubation, the gels were rinsed briefly with water and stained with 0.02% Ruthenium Red (Sigma) for 1h at room temperature. The dye was then poured off and the gel was rinsed with water again. Diameters of the red halos around each well were measured using the Fiji software.

### Mass spectrometry

Each SDS-PAGE band was manually excised from the gels to be hydrolysed according to (Shevchenko et al. 1996). Each sample was destained in a 40% ethanol-10% acetic acid solution (v/v) for 15 min and progressively dehydrated with 15 min incubations in first a 50% acetonitrile-25 mM ammonium bicarbonate solution and then in 100% acetonitrile. The dehydrated gel pieces were then reduced for 30 min with a 10 mM solution of DTT at 56°C and alkylated for 45 min with 55 mM iodoacetamide. The samples were then dehydrated as above. For digestion, the samples were incubated overnight at 37°C with 0.01 μg/μL of trypsin (Promega, Madison, WI) and dissolved in 50 mM ammonium bicarbonate, pH 8. The tryptic fragments were extracted from the gel by centrifugation after incubation of 15 min in a 0.5% trifluoroacetic acid - 50% acetonitrile (v/v) and in 100% acetonitrile solution, dried in a speed-vac, and reconstituted with 25 μL of a 2% acetonitrile - 0.1% formic acid (v/v) buffer. HPLC was performed on a NanoLC-Ultra system (Eksigent). A 4 μL sample of the peptide solution was loaded at 7.5 μL min^−1^ on a precolumn cartridge (stationary phase: C18 Biosphere, 5 μm; column: 100 μm inner diameter, 2 cm; Nanoseparations) and desalted with 0.1% fluorhydric acid and 2% acetonitrile. After 3 min, the precolumn cartridge was connected to the separating PepMap C18 column (stationary phase: C18 Biosphere, 3 μm; column: 75 μm inner diameter, 150 mm; Nanoseparations). Buffers A and B respectively were prepared with 0.1% HCOOH in water, and with 0.1% HCOOH in acetonitrile. The peptide separation was achieved at 300 nL min−1 with a linear gradient: from minute 0 to 10, at 5 to 30% of B; from minute 10 to 12, at 30 to 95% of B; from minute 12 to 18, at 95% of B (the regeneration step); from minute 18 to 19, at 95 to 5% of B; from minute 19 to 20, at 5% of B (the equilibration step). Remaining percentages are those of buffer A. One run took 45 min. Eluted peptides were online analyzed with an LTQ XL ion trap (Thermo Electron) using a nanoelectrospray interface. Ionization (1.5 kV ionization potential) was performed with liquid junction and a non-coated capillary probe (10 μm inner diameter; New Objective). Peptide ions were analyzed using Xcalibur 2.07 with the following data-dependent acquisition steps: (1) full MS scan (mass-to-charge ratio (m/z) 350-1 400, centroid mode) and (2) MS/MS (isolation width 1m/z, qz = 0.25, activation time = 30 ms, and normalized collision energy = 35%; centroid mode). Step 2 was repeated for the three major ions detected in step 1. Dynamic exclusion was set to 45s.

A database search was performed with XTandem (version 10-12-01-1) (http://www.thegpm.org/TANDEM/). Enzymatic cleavage was defined as a trypsin digestion with one possible miscleavage. Regarding protein modifications, Cys carboxyamidomethylation was set to static. As potential modifications, Met oxidation, Nter acetylation, Nter deamidation on Gln, Nter deamidation on carbamidomethylated Cys, Nter neutral loss on Glu were set. Precursor mass and fragment mass tolerance were 2.0 and 0.8, respectively. A refinement search was added with similar parameters. The Araport11_genes.201606 database and a contaminant database (trypsin, keratins) were used for peptides identification. Identified peptides were filtered and grouped using Xtandem according to: (1) A minimum of two different peptides required with an E-value smaller than 0.05, (2) a protein log(E-value) (calculated as the product of unique peptide E-values) smaller than -4.

### Growth inhibition assays

Arabidopsis seedlings were grown for three days on vertical petridishes on ½ MS medium (Duchefa, reference M0221) with 1.2% phytoagar (Duchefa, reference: P1003) at 22 °C, 60% HR, 50 μmol.m^−2^.s^−1^ white light (light source LED) with 18h-8h day-night cycle, before transfer to microtiter plates (4 seedlings per well) containing 100 μl of liquid ½ MS medium supplemented or not with PMEI3 or the PMEI inhibitor epigallocatechin-3-gallate (EGCG) as a positive control. The root length was scored 14h later. Statistics were performed using the R statistics platforms. Significance (α = 0.05) was assessed by analysis of variance (ANOVA ; parametric) using linear mixed-effect models followed by Tukey HSD.

### COS^647^ labeling of roots

Seedlings were fixed for 10 min in 10% acetic acid and 3.8% formaldehyde in “2F4 buffer” (Haas et al., 2020) and neutralized overnight in 10mM NH^4^Cl in “2F4 buffer”. After washing with 25mM MES buffer pH5.7, the samples were incubated in the same buffer for 10 min with 1/500 COS^647^ (Mravec et al., 2014) and, following washing, imaged with a confocal (Zeiss 710). COS^647^ was kindly provided by Jozef Mravec (University of Copenhagen, DK).

## Acknowledgements

Gladys Cloarec is thanked for assistance in the use of the confocal microscope. The work was financed by ANR projects “Pectosign” and “Homeowall” and the CSC PhD fellowship Nr 201506300059 to FX. The IJPB benefits from the support of Saclay Plant Sciences-SPS (ANR-17-EUR-0007). This work has benefited from the support of IJPB’s Plant Observatory technological platforms.

## Supplementary data and Figures

**Plant codon-optimized *PMEI3* DNA sequence** CTGCAGGAAGACATAACGGTGCTGAGGATATCGTTCACTCTTCTTGTGAGCACGCTTCTTACCCATCCTTGTGTGTTAGAACCTTGTCCTCTTACTCCGGTCCAACCATCACTAACAGAAGAGACTTGGCTCAGGCCGCTATCAAGATTTCTTTGTCTCACGCTCAATCCGCCGCTAAGAAGTTGGCTGTTGTTAGAGACTCCGTCGGTAAGAAGAAGCAAGAGAAGGCTGCTTTGGTTGACTGCGTTGAGATGATTGGTGACTCCGTTGACGAGTTGTCCAGAACCTTGGGTGTTTTGAAGCACTTGAGAGTTTCTGGTGGTTCCGCCAAAGAATTCAGATGGCAAATGTCCAACGCTCAGACTTGGGCTTCTGCTGCTTTGACTGATGACGACACTTGTCTGGATGGTTTCCAAGGTATGGACGACGGTGAGATCAAGACTGAGGTTAAGCAGTGGATGACCAAGGTTGCTAGAGTTACTTCCAACGCCTTGTACATGGTCAACCAGTTGGACGAGACTAGAGGTAAGCCACACGACGTTCACTTGTT TCTAGA

**Figure S1: in vitro activity of PME preparations used in this study**. Error bars are SD (n=3)

**Figure S2: Purification of PMEI3 (A) and Gel diffusion assay for PME and PMEI activity**. A: Coomassie-stained gel of the affinity purification of PMEI3, with 3 successive elutions from the affinity column.

**Figure S3: PME3 RNA expression pattern in *Arabidopsis thaliana***. Source plant eFP viewer on TAIR (Klepikova et al., 2016).

**Figure S4: PME2 RNA expression pattern in *Arabidopsis thaliana***. Source plant eFP viewer on TAIR (Klepikova et al., 2016).

**Figure S5: Cell type-specific RNA expression pattern of PME2 and PME3 in *Arabidopis thaliana* roots**. Source: https://bioit3.irc.ugent.be/plant-sc-atlas/ (Wendrich et al., 2020)

**Figure S6: *Root growth inhibition by PME inhibitor EGCG***. Representative photographs (A-G) and primary root lengths (H) of 5-day-old *Arabidopsis thaliana* treated with increasing concentrations of PME inhibitor EGCG (Numbers are in μM). Error bars are SD (n > 30).

## Notes

### Competing Interest Statement

The authors have declared no competing interest.

