## Supplementary material for "Biochemical characterization of Pectin Methylesterase Inhibitor 3 from *Arabidopsis thaliana*": Xu et al Suppl Figures

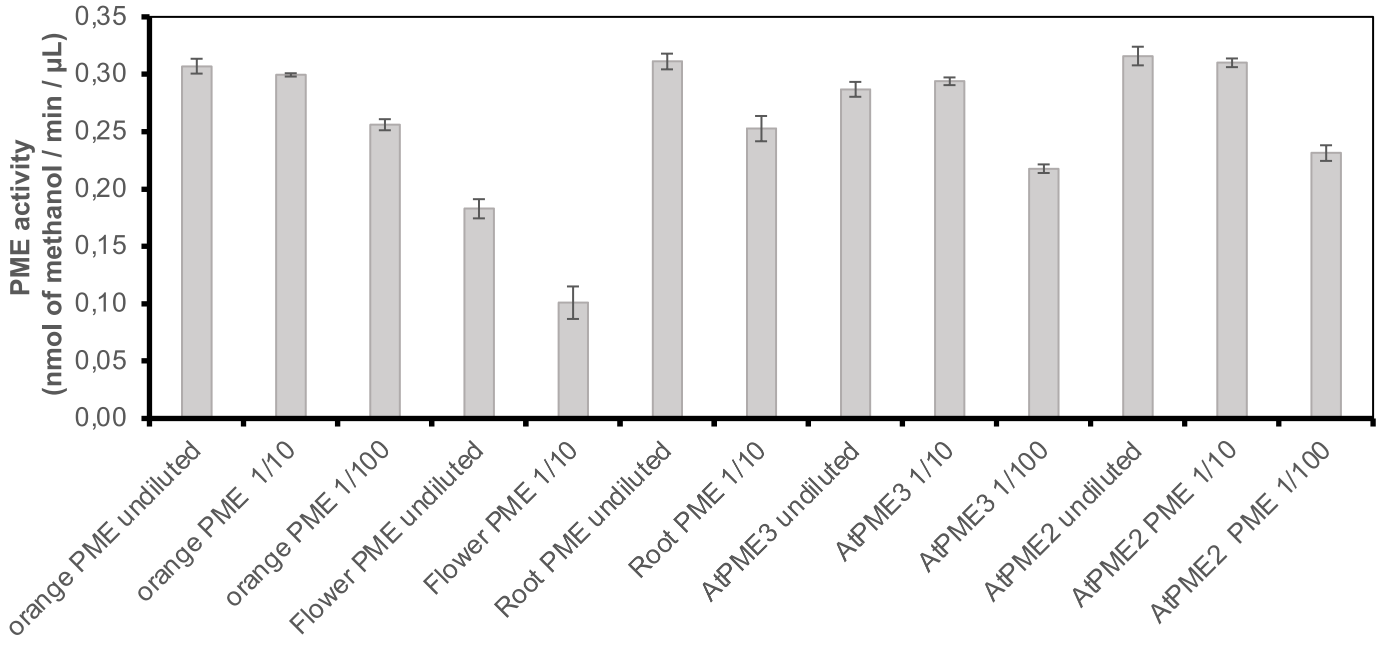


**Figure S1: in vitro activity of PME preparations used in this study.**
Error bars are SD (n=3)

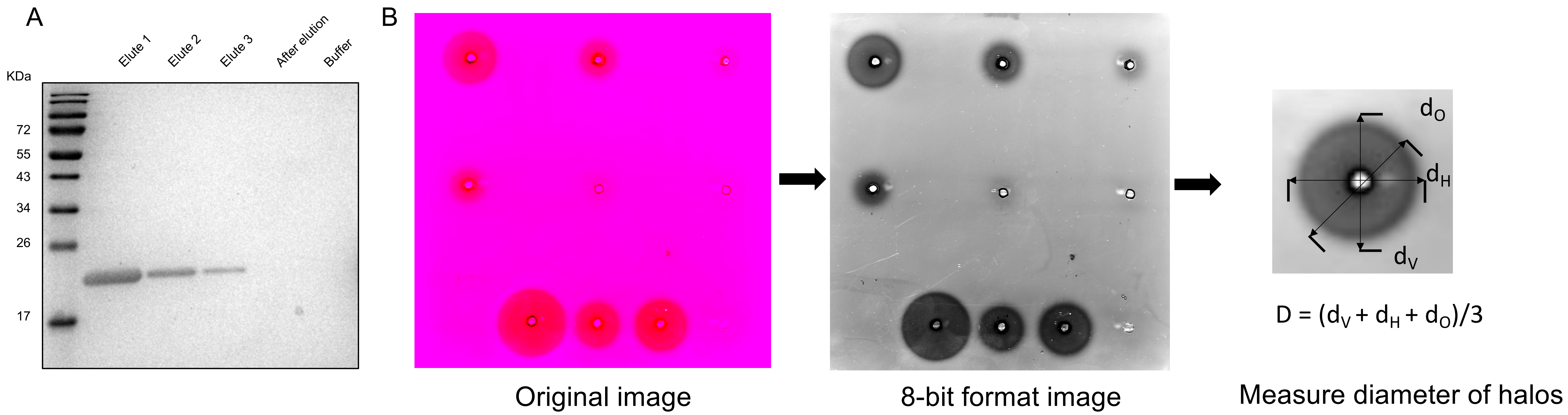


**Figure S2: Purification of PMEI3 (A) and Gel diffusion assay for PME and PMEI activity. A:** Coomassie-stained gel of the affinity purification of PMEI3, with 3 successive elutions from the affinity column.

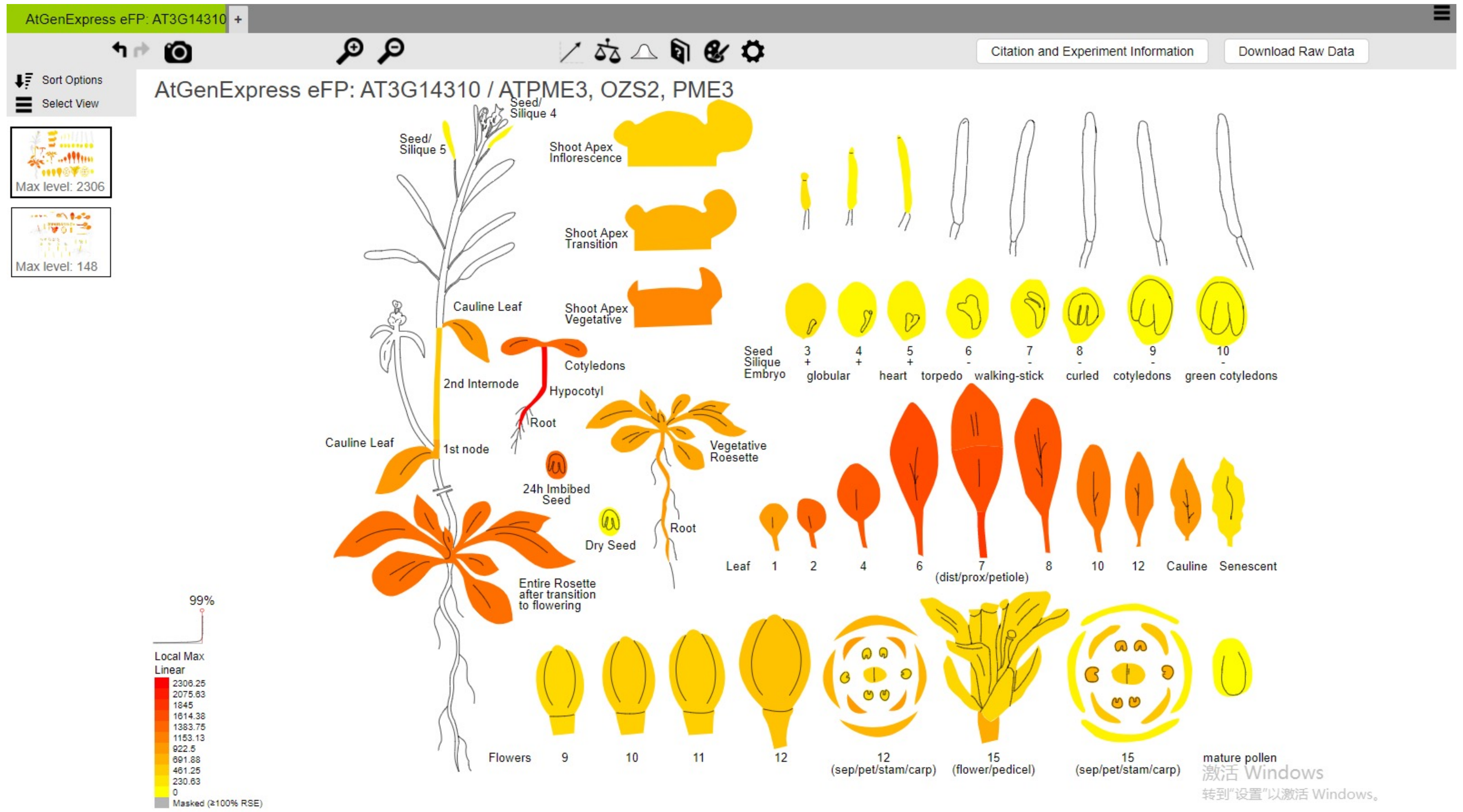


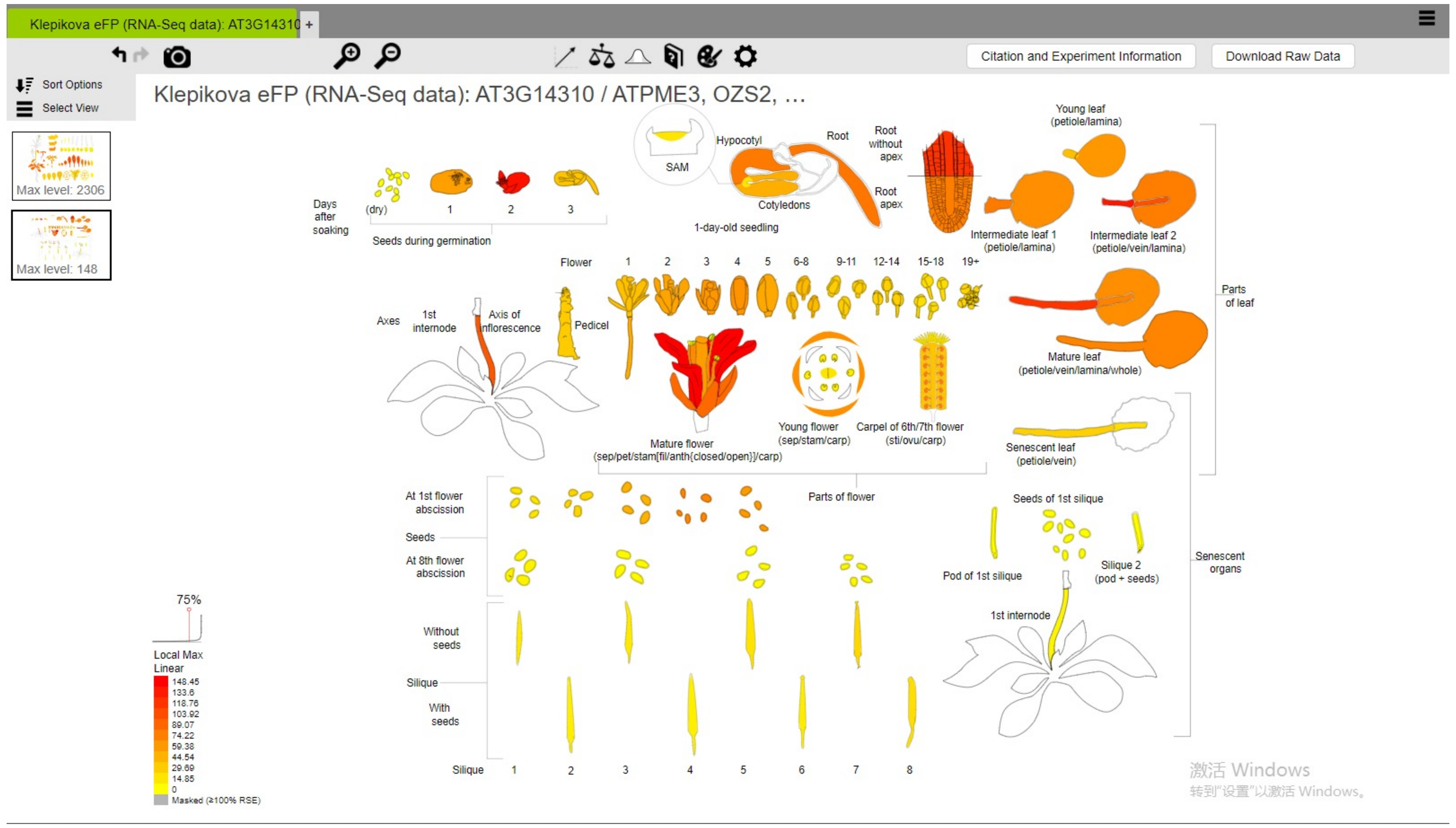


**Figure S3: PME3 RNA expression pattern in *Arabidopsis thaliana*.** Source plant eFP viewer on TAIR (Klepikova et al., 2016)

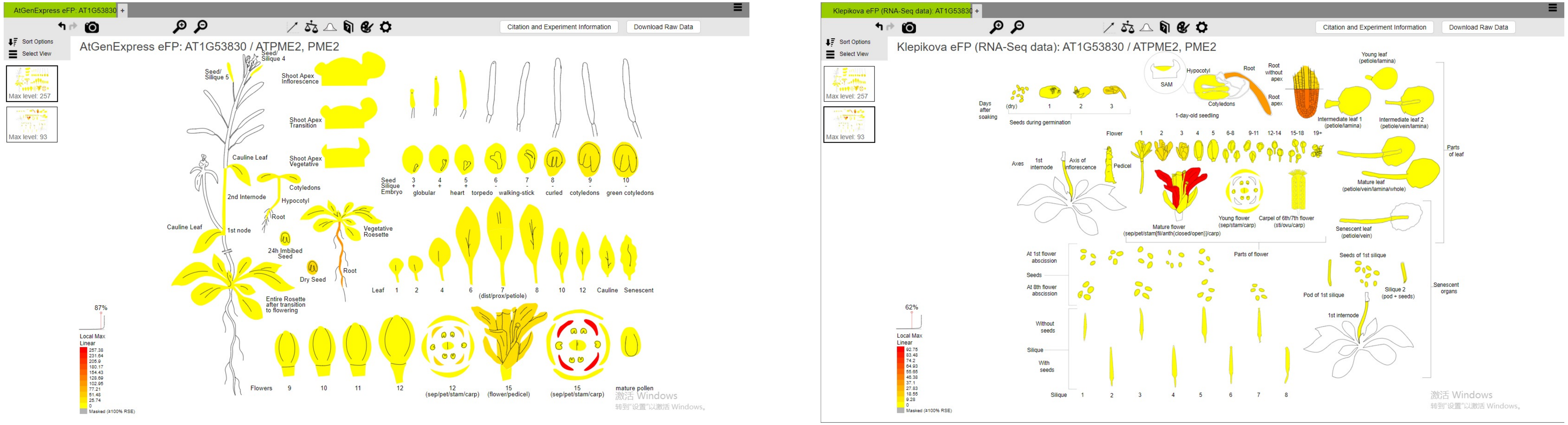


**Figure S4: PME2 RNA expression pattern in *Arabidopsis thaliana*.** Source plant eFP viewer on TAIR (Klepikova et al., 2016).

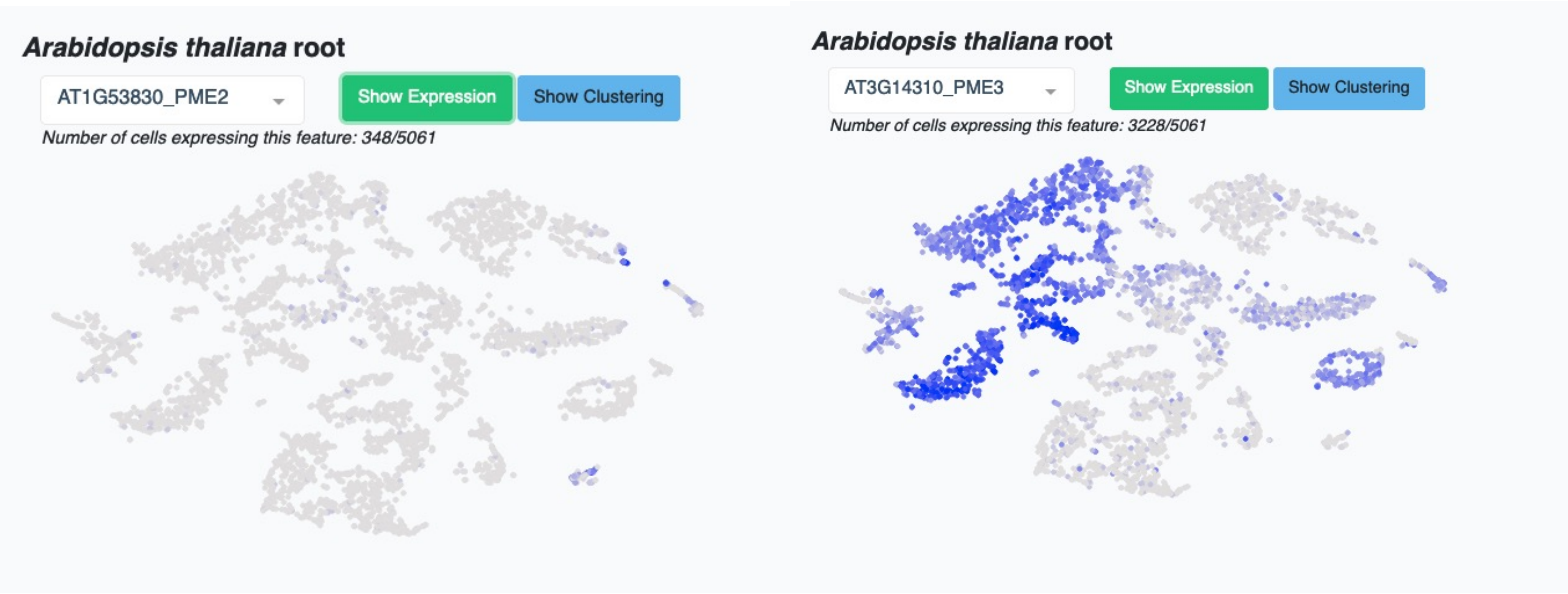


https://bioit3.irc.ugent.be/plant-sc-atlas/

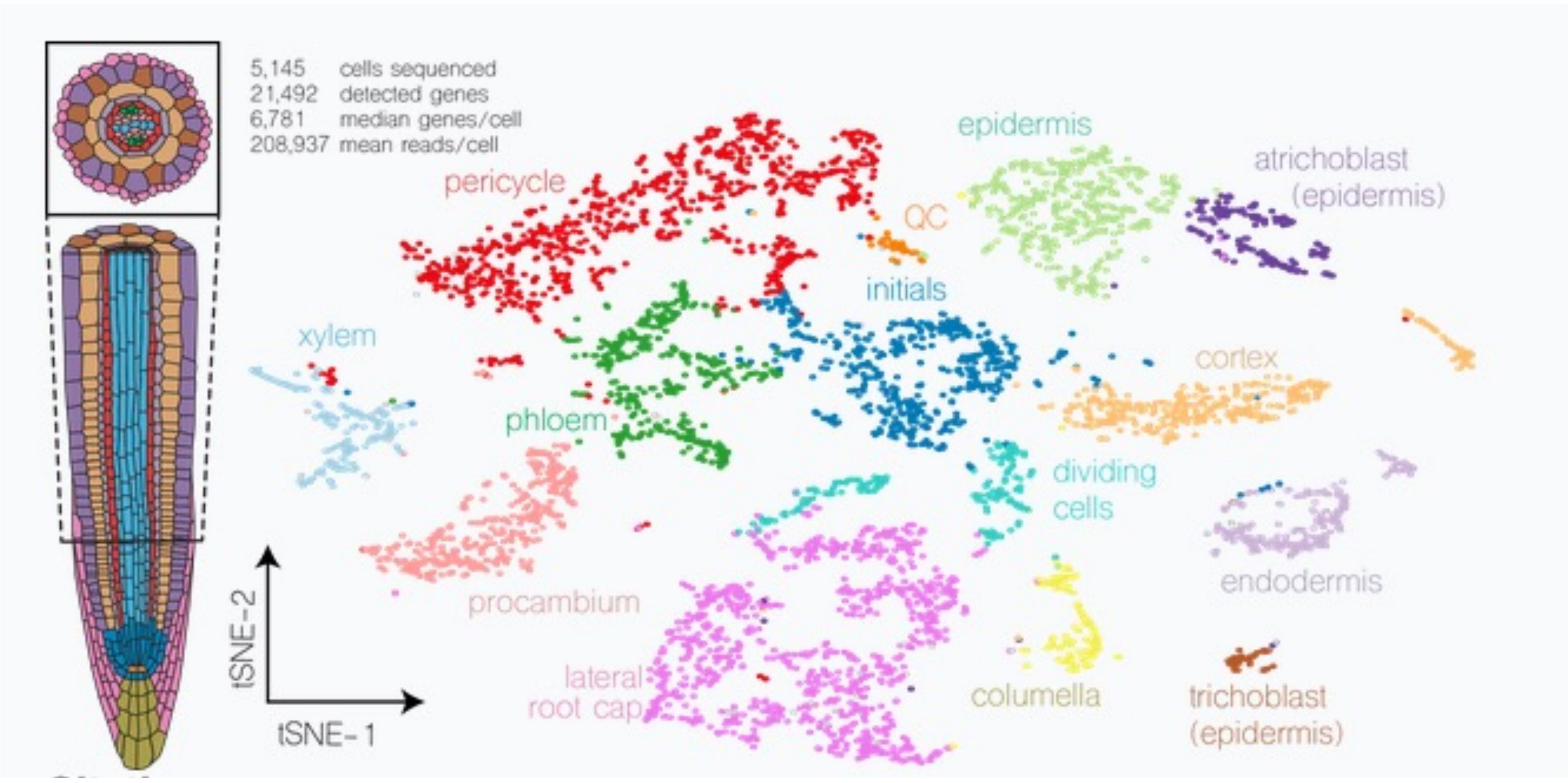


**Figure S5: Cell type-specific RNA expression pattern of PME2 and PME3 in *Arabidopsis thaliana* roots.** Source: https://bioit3.irc.ugent.be/plant-sc-atlas/ (Wendrich et al., 2020)

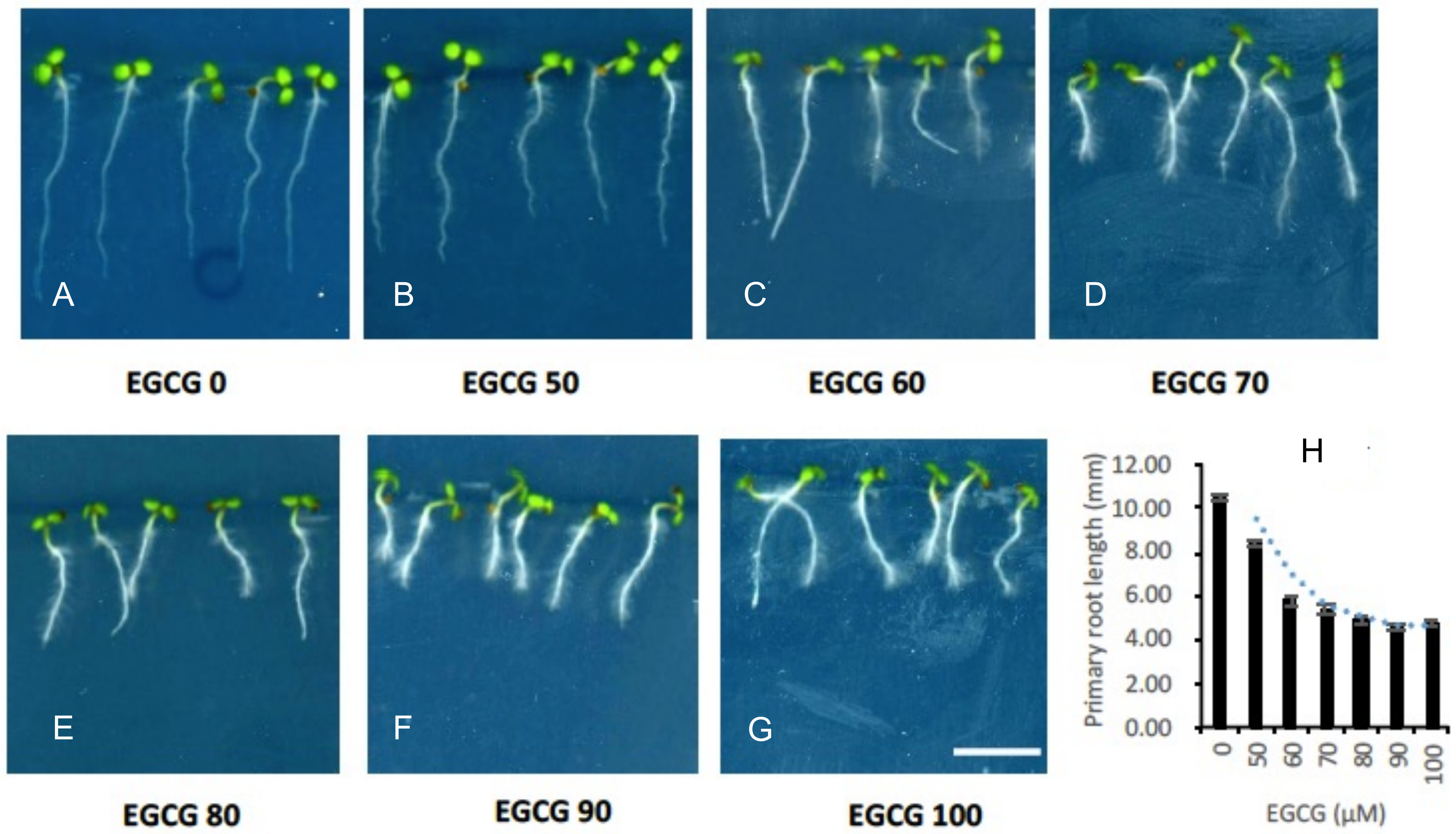


**Figure S6: *Root growth inhibition by PME inhibitor EGCG*.** Photographs (A-G) and primary root lengths (H) of 5-day-old *Arabidopsis thaliana* treated with increasing concentrations of PME inhibitor EGCG (Numbers are in µM). Error bars are SD (n > 30).
